# JNK signalling regulates antioxidant responses in neurons

**DOI:** 10.1101/2020.06.05.136622

**Authors:** Chris Ugbode, Nathan Garnham, Laura Fort-Aznar, Gareth Evans, Sangeeta Chawla, Sean T. Sweeney

## Abstract

Reactive oxygen species (ROS) are generated during physiological bouts of synaptic activity and as a consequence of pathological conditions in the central nervous system. How neurons respond to and distinguish between ROS in these different contexts is currently unknown. In Drosophila mutants with enhanced JNK activity, lower levels of ROS are observed and these animals are resistant to both changes in ROS and changes in synapse morphology induced by oxidative stress. In wild type flies, disrupting JNK-AP-1 signalling perturbs redox homeostasis suggesting JNK activity positively regulates neuronal antioxidant defense. We validated this hypothesis in mammalian neurons, finding that JNK activity regulates the expression of the antioxidant gene *Srxn-1*, in a c-Jun dependent manner. We describe a conserved ‘adaptive’ role for neuronal JNK in the maintenance of redox homeostasis that is relevant to several neurodegenerative diseases.

## Introduction

Active neurons generate reactive oxygen species (ROS) predominantly as a by-product of mitochondrial respiration. ROS levels are neutralized by constitutive and adaptive reductive mechanisms operating in neurons and glia, including the glutathione system [1–3]. In this manner, the amplitude and temporal dynamics of the ROS signal are controlled, damage is limited and transient ROS signals can be interpreted in part to support the growth and plasticity of neurons [4, 5]. A physiological level of ROS has been demonstrated to regulate a range of nervous system processes, including neuronal development, synaptic plasticity, and neural circuit tuning [5–7].

In many neurodegenerative disorders the reductive capacity of neurons is overwhelmed, contributing to disease progression [4, 8]. These damaging levels of ROS, termed oxidative stress, overwhelm neuronal antioxidant defenses. A central component of the neuronal response to ROS is the activation of c-Jun N-terminal kinase, JNK. ROS have been shown to activate JNK-AP-1 signalling [9] which regulates neuronal growth and plasticity [10–13]. In *Drosophila* models, excessive ROS driving increases in synaptic growth are seen in both activity generated excitotoxicity and lysosomal storage disease [5, 12, 14]. Similar changes in synaptic structure can be induced directly by application of oxidants such as paraquat [12] or diethylmaleate (DEM) [5], or through genetic activation of JNK via manipulation of the JNKK *hemipterous [15]* (*hep*) or the JNKKK *Wallenda [11]* (*wnd*), upstream activators of JNK.

Whether JNK activation by ROS in neurons is protective or degenerative is unknown. Recent evidence indicates JNK-AP-1 activity can prevent degeneration of injured axons [16] while JNK inhibition prevents neuronal loss after injury by promoting AKT signaling [17].

We recently found that knockdown of enzymes involved in antioxidant defense, superoxide dismutase and catalase, reshapes neuronal morphology at the *Drosophila* larval neuromuscular junction (NMJ) [5]. Similarly, mis-expressing antioxidant enzymes to manipulate levels of individual ROS species has profound effects on synaptic and dendritic growth and arborisation [5, 12]. This suggests that plasticity at the synapse is finely tuned to the level of ROS. Given that JNK is known to regulate synaptic plasticity in response to ROS and that knockdown of antioxidant genes can change synapse structure, we hypothesized that JNK-induced structural changes in neurons are mediated in part by an antioxidant response.

To understand if JNK activity regulates antioxidant responses in neurons, we monitored ROS levels in *Drosophila* with mutations in *highwire* (*hiw*) and *puckered* (*puc^E69^*). In *hiw* mutants, unrestrained *wallenda* (*wnd*) activity activates JNK and downstream AP-1 signaling to generate exuberant synaptic overgrowth [11, 18]. *Puc* encodes a dual specificity phosphatase, transcriptionally activated by AP-1 that counteracts JNK activity [19]. In both the *hiw* and *puc^E69^/+* mutant backgrounds we identified enhanced antioxidant defenses and low levels of ROS. In *hiw*, this enhanced antioxidant defense can be reversed in neurons by down-regulating JNK-AP-1 signaling. Conversely, both *puc^E69^*/+ and *hiw* flies are resistant to chemically induced oxidative stress, while inhibition of JNK-AP-1 signaling increases ROS levels in wild type neurons. We show that in mammalian neurons exposed to oxidative stress, the JNK-AP-1 pathway coordinates the transcription and translation of the antioxidant gene *Sulfiredoxin-1*, which was localised predominantly in synaptic compartments. Our data indicates that activation of JNK in neurons drives antioxidant responses to shape synaptic plasticity.

## Results

### JNK Activity Drives Antioxidant Responses

To quantify levels of ROS in *Drosophila*, we used the Amplex red assay for detection of hydrogen peroxide [20]. We have previously found that food containing 5mM and 10mM DEM which depletes glutathione levels, is sufficient to induce an overgrowth at the larval NMJ [5], with 10mM concentrations being detrimental to survival (Fig S1, 6% of flies pupate). To understand whether DEM influences ROS levels in flies, we raised flies on food containing ethanol (vehicle) or 5mM DEM. Using Amplex red we found that wild type flies have an increased ROS burden when raised on food containing 5mM DEM (Fig 1A). To investigate how JNK activity influences ROS levels under these conditions, we reared *puc^E69^*/+ and *hiw* mutants on the same food and found that both genotypes have significantly lower levels of ROS than wild type flies in vehicle-containing food. Moreover, both genotypes failed to show an increase in ROS when reared on DEM-containing food (Fig 1A) suggesting that they are resistant to the oxidative stress generated by DEM.

**Figure 1:**
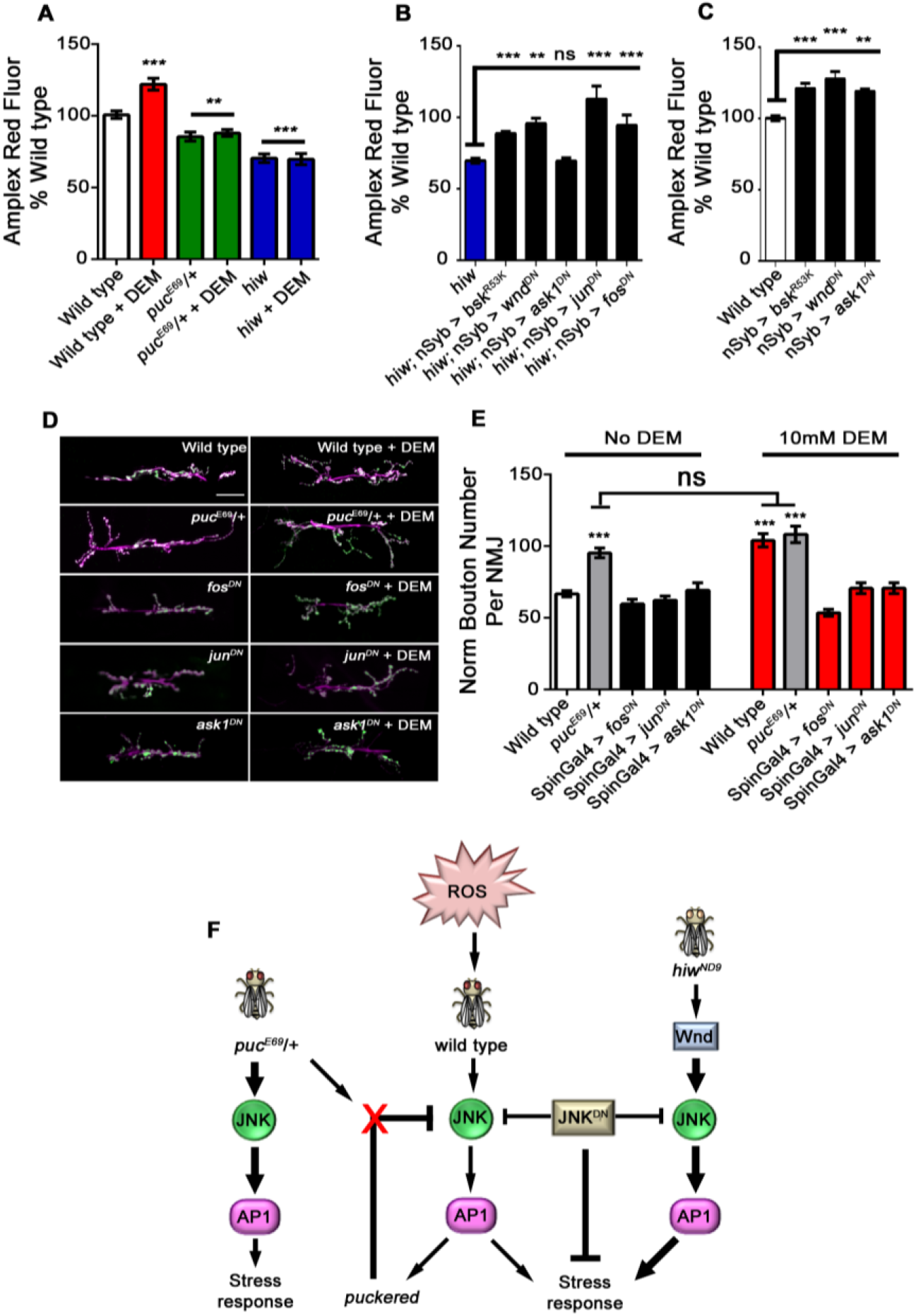
JNK activity regulates neuronal antioxidant responses. **A** DEM induces oxidative stress in wild type flies but not *puckered* (*puc^E69^*/+) or *highwire* (*hiw*) mutants. Quantification of hydrogen peroxide levels (Amplex red fluorescence) in wild type, *puc^E69^*/+ or *hiw* mutant flies administered veh (0.16% ethanol) or DEM (5mM). **B** Quantification of hydrogen peroxide levels in *hiw* mutants and *hiw* mutants pan-neuronally expressing dominant negative *jnk* (*h*; nSyb > *bsk^R53K^*), *wnd* (*h*; nSyb > *wnd^DN^*), *ask1* (*h*; nSyb > *ask1^DN^*), *jun* (*h*; nSyb > *jun^DN^*) and *fos* (*h*; nSyb > *fos^DN^*). **C** Quantification of hydrogen peroxide levels in wild type flies, pan-neuronally expressing dominant negative *jnk* (nSyb > *bsk^R53K^*), *wnd* (nSyb > *wnd^DN^*) and *ask1* (nSyb > *ask1^DN^*). **A-C** Graphs show amplex red fluorescence values normalized to the average signal of wild type flies. Data are plotted as mean ± SEM (** p < 0.01; *** p < 0.001; one-way ANOVA; minimum 15 flies per genotype). **D** Representative micrographs showing synaptic overgrowth at the *Drosophila* third instar larval NMJ (Muscle 6/7, hemi-segment A3) in wild type, *puc^E69^*/+ mutants and wild type flies expressing dominant negative *fos* (*fos^DN^*), *jun* (*jun^DN^*) and *ask1* (*ask1^DN^*), reared on food containing ethanol or DEM. Scale bar = 30 μm. **E** Quantification of mean normalized bouton number from (**D**) genotypes. Data are plotted as mean ± SEM (*** p < 0.001; one-way ANOVA; minimum 15 NMJ’s analysed per genotype). **F** Schematic showing the contribution of JNK to neuronal antioxidant stress responses in *Drosophila*.

Given that both *puc^E69^*/+ and *hiw* mutants have lower levels of ROS than wild type flies and are resistant to chemically-induced oxidative stress, we hypothesized that if JNK activity could be reduced in neurons of these flies, then ROS levels would increase back to wild type. To test this, we used *hiw* mutants and expressed dominant negative transgenes of *jnk*, *wnd*, *ask1*, *jun* and *fos* pan-neuronally using nSyb-Gal4 (Fig 1B). We found that pan-neuronal expression of these dominant negative transgenes increased ROS levels of *hiw* mutants back to wild type controls, except for *ask1^DN^*. This data indicates that manipulation of JNK, its upstream regulators and downstream targets, regulates ROS levels in neurons.

To further evaluate the relationship between JNK activity and ROS levels, we investigated whether expression of dominant negative *jnk, ask1 and wnd* affects ROS levels in wild type flies. We found that expression of all dominant negative transgenes, pan-neuronally, significantly increases the ROS levels of wild type flies (Fig 1C). Indeed, the levels observed were similar to those observed when wild type flies are reared on food containing DEM.

The elaborate synaptic morphology of *hiw* mutant flies has been extensively described [11] and is a consequence of activation of the JNK-AP-1 pathway. We therefore chose to evaluate the synaptic morphology of *puc^E69^*/+ neurons by analyzing the NMJ at Muscle 6/7, hemi-segment A3. Similar to *hiw* mutants, we found that *puc^E69^*/+ larvae have overgrown NMJ synapses (Fig 1D). DEM feeding also causes an overgrowth of the NMJ when compared to wild type flies reared on normal food. Feeding of DEM has no effect on the NMJ of *puc^E69^*/+ larvae, indicating that JNK activity alone is sufficient to change synaptic morphology, which correlates with an antioxidant response (decrease in ROS levels from Fig 1A). Furthermore, the DEM induced changes in synaptic plasticity in wild type flies, can be rescued when dominant negative transgenes of *jun, fos* and *ask1* are expressed pre- and post-synaptically using *spin*GAL4 (Figure 1D+E). Our observations in *Drosophila* show that JNK activity regulates synaptic morphology but also antioxidant responses. In *puc^E69^*/+ and *hiw* flies, JNK-AP-1 signalling generates an antioxidant response which lowers the total levels of ROS in flies and protects them from induced oxidative stress while also causing an increase in NMJ synapse size. In wild type flies, inducing oxidative stress changes synaptic morphology, generates a stress response and these responses can be regulated by JNK (Figure 1F).

### DEM induces oxidative stress in primary mammalian neurons

JNK activity in mammalian neurons is often associated with apoptosis and activated JNK is commonly used as a hallmark of cell death in neurodegenerative disease. Given that JNK-AP-1 signalling in the fly generates an antioxidant response and decreasing JNK activity in fly neurons increases ROS levels, we hypothesized that antioxidant responses in mammalian neurons would also be regulated by JNK-AP-1. Having previously validated the effects of DEM in the fly, we established a mammalian model of oxidative stress, using DEM to deplete cellular glutathione. We used this model to assay antioxidant responses after DEM treatment. We first characterized the effect of DEM on cellular GSH levels and ROS production in mature primary rat neurons, using a glutathione assay measuring total glutathione (oxidized and reduced forms). Figure S2 A shows that application of DEM to primary cortical neuronal cultures caused a concentration and time-dependent decrease in cellular glutathione. A 1 h and 4 h treatment of DEM at concentrations of 10 μM and 100 μM caused a rapid decrease in GSH (Fig S2A). Lower concentrations of 1 μM DEM reduced cellular GSH content with less severity. Given the extent of GSH depletion by DEM, we investigated whether DEM induces mitochondrial toxicity by monitoring WST-1 absorbance (Water Soluble Tetrazolium Salts) as a measure of mitochondrial stress. Compared to vehicle treated neurons, DEM induced a small but significant decrease in mitochondrial capacity, however this was much less than treatment with equivalent concentrations of paraquat (PQ) and rotenone (RT) (Fig S2B). To determine if DEM induces a ROS burden as it does in our fly model we measured H_2_O_2_ levels in neurons treated with DEM using Amplex red reagent. The Amplex red assay revealed that DEM (100 μM) significantly increased in H_2_O_2_ levels (Fig. S2C), after 24 hours compared to ethanol treated controls (Fig. S2C). DEM induced oxidative stress is significantly attenuated when neurons are pre-incubated with catalase. The duration and concentrations of DEM that we employed deplete glutathione and induce an increase in ROS with minimal toxicity to mitochondrial function.

To model chronic oxidative stress, we treated neurons for longer time points (48 hr) with DEM (100μM) and found that DEM causes a retraction of the dendritic arbor. This effect is rescued when neurons are pre-incubated with Catalase (100nM, Fig S2D) and when the components that produce glutathione, the catalytic and modifying subunits of glutamate cysteine ligase (GCL), are transduced into neurons (Fig S3).

### Oxidative stress activates JNK-dependent SRXN-1 expression

JNK enzymes are key mediators of cellular responses to stress in neurons [21], including oxidative stress [22, 23], and alterations in JNK activity have previously been linked to changes in dendritic complexity [24]. Phosphorylation of threonine 183 and tyrosine 185 increase JNK activity [21]. To characterize the effectiveness of both DEM and the JNK inhibitor SU 3327 on regulating JNK phosphorylation and activation, we treated primary neurons with DEM (100μM, 1 hr) alone or in the presence of SU 3327 (700nM, 1 hr pre-treatment) and assayed JNK phosphorylation (Fig 2A). We found a significant increase in phospho-JNK/total JNK ratios when neurons were treated with DEM, which was significantly attenuated when JNK activity was blocked with SU 3327 (Fig 2 A+B).

**Figure 2:**
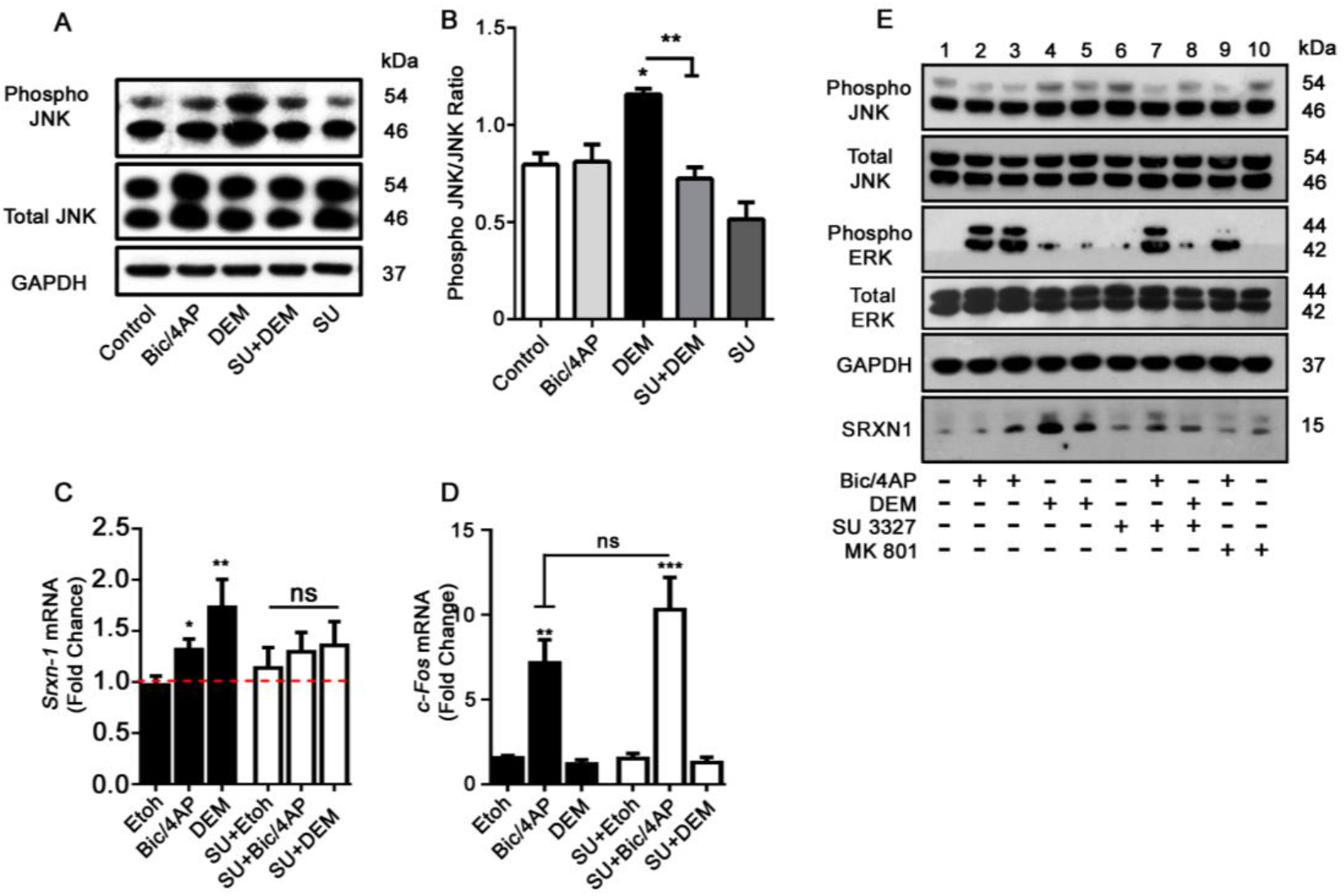
ROS induced JNK phosphorylation triggers neuronal antioxidant responses. ***A*** Representative western blot of neurons treated with Bicuculine (50 μM, Bic) and 4-Aminopyridine (500 μM, 4AP) to increase synaptic activity (Bic/4AP) or DEM (100μM) for 1 hour, to induce oxidative stress. JNK activity was inhibited by a 1 hour pre-treatment with SU-3327 (SU, 700nM). ***B*** Quantification of total phospho-JNK/pan-JNK ratios (* p < 0.05; ** p < 0.01; one-way ANOVA; n = 3 biological replicates). ***C+D*** The mRNA levels of *Srxn-1* and *c-Fos* were determined using quantitative PCR (qPCR) and were normalized to *Gapdh* mRNA levels. mRNA levels are expressed relative to untreated controls and shown as means ± SEM (* p < 0.05; ** p < 0.01; unpaired t-Test; n = 4 biological replicates) ***C*** *Srxn-1* mRNA is induced by oxidative stress in a JNK dependent manner. ***D*** JNK signaling downstream of oxidative stress does not require c-Fos. ***E*** JNK regulates SRXN-1 expression under oxidative stress but not during synaptic activity. Representative western blot probed for SRXN-1, Pan-JNK, phospho-JNK, Pan-ERK, phospho-ERK and GAPDH antibodies. Neurons were treated with Bic/4AP (50/500 μM) or DEM (100 μM) in the presence or absence of either the JNK inhibitor SU 3327 (700nM) or the NMDA receptor blocker MK801 (10μM) for 30 mins (lanes 2/4/6) or 4 h (lanes 3/5/7/8/9).

Bicuculline, a competitive antagonist of GABA_A_ (γ-aminobutyric acid) receptors, and 4-aminopyridine, a voltage activated potassium blocker, are commonly used pharmacological tools that increase synaptic activity in cultures of cortical neurons [25]. These tools have shown that synaptic activity can upregulate neuronal antioxidant genes, in an ERK-AP-1 dependent manner. A putative antioxidant target gene of this signaling pathway, is *Sulfiredoxin-1* (*Srxn-1)*[26]. To identify whether JNK-AP-1 signalling could regulate neuronal antioxidant genes, we compared our experiments with neurons treated with Bicuculline (Bic, 50μM) and 4-Aminopyridine (4-AP, 500μM) as a positive control (Bic/4AP).

Using DEM, we found that JNK activity regulates *Srxn-1* transcription. DEM (10 μM) induced a 1.73-fold induction in *Srxn-1* mRNA expression compared to the ethanol treated controls (Fig. 2C). When neurons were pre-treated with the JNK inhibitor SU 3327 for 24 h, DEM failed to induce *Srxn-1* mRNA. Similar to DEM treatment, increasing synaptic activity with Bic/4AP increased *Srxn-1* mRNA (1.32-fold), however this induction was unaffected by JNK inhibition.

Synaptic activity is known to induce expression of *c-Fos*, an AP-1 component. We determined whether oxidative stress also induced *c-Fos* transcription. Figure 2D shows that there was no change in *c-Fos* mRNA levels with DEM treatment whereas Bic/4AP treatment robustly induced *c-Fos* transcription. Similar to activity-induced *Srxn-1* induction, *c-Fos* induction by synaptic activity was unaffected by inhibition of JNK (Fig. 2D) suggesting that synaptic activity and oxidative stress recruit signaling pathways differentially to activate potentially divergent gene transcription responses. We next evaluated whether JNK activity regulates SRXN-1 protein expression. Figure 2E shows that treatment of neurons with DEM (100 μM) rapidly increased SRXN-1 protein expression within 30 min, with high levels detected at 4h as assessed by western blot using an antibody that we first validated by overexpressing human SRXN-1 in HEK293T cells (Fig S4 A). DEM-induced SRXN-1 expression was attenuated when neurons were pre-treated with SU 3327 (700 nM, 1 h pre-incubation). As expected, and in agreement with previous work [26], synaptic activity also induced SRXN-1 expression after a 4 h treatment with Bic/4AP, which was attenuated when neurons were pre-treated for 1 h with the non-competitive NMDA receptor antagonist MK801 (10 μM) but was unaffected by JNK inhibition. Moreover, synaptic activity but not DEM, induced ERK activation (Figure 2E). Taken together, these data indicate that oxidative stress activates JNK to coordinate an antioxidant response in neurons, which is distinct from antioxidant responses coordinated by synaptic activity.

### JNK-c-Jun signaling regulates antioxidant responses

To identify the cellular mediators that co-ordinate SRXN-1 expression under conditions of synaptic activity or oxidative stress, we assessed the expression and interaction of putative AP-1 components under the two conditions. Cortical neurons were treated with Bic/4AP or DEM for 1 or 4 h and c-Fos or c-Jun immunofluorescence was quantified in NeuN positive neurons. In neurons treated with DEM (Figure 3A and 3B) c-Fos immunofluorescence was found to decrease. In contrast, DEM induced a significant increase in c-Jun immunofluorescence. As expected, in neurons stimulated with Bic/4AP for 1 or 4 h, both c-Fos and c-Jun immunofluorescence was found to increase (Fig S4, B+C). These data suggest that oxidative stress relies on Jun proteins to mount an antioxidant response in the absence of c-Fos. To further investigate the differential involvement of specific AP-1 components we immunoprecipitated c-Jun from cortical neurons treated for 1 h with either DEM or Bic/4AP and assessed its phosphorylation and association with c-Fos protein. While both DEM and Bic/4AP induce c-Jun expression and phosphorylation, c-Jun co-immunoprecipitated with c-Fos following Bic/4AP-induced synaptic activity but not during oxidative stress mediated by DEM (Fig 3C). In mammals, c-Jun can form heterodimers with other members of the Jun family to form the dimeric AP-1 transcription factor. To identify if c-Jun was associated with another Jun family member, we probed membranes for JunB and JunD and found no association between either protein with c-Jun in DEM-treated neurons (Fig S4, D). To determine whether c-Jun alone is sufficient to induce SRXN-1 expression, lentiviral particles encoding human c-Jun (Fig 3D) were transduced into cortical neurons. Lentiviral c-Jun expression was sufficient to induce SRXN-1 expression in neurons (Fig 3E), however activation of Nuclear Factor Erythroid 2-related factor 2 (Nrf2) using the well characterized agonist TBHQ (10μM, 16 h) had no effect on neuronal expression of SRXN-1. Taken together, these data suggest that oxidative stress activates JNK, which selectively recruits c-Jun to induce SRXN-1 expression in neurons (Fig 3F).

**Figure 3:**
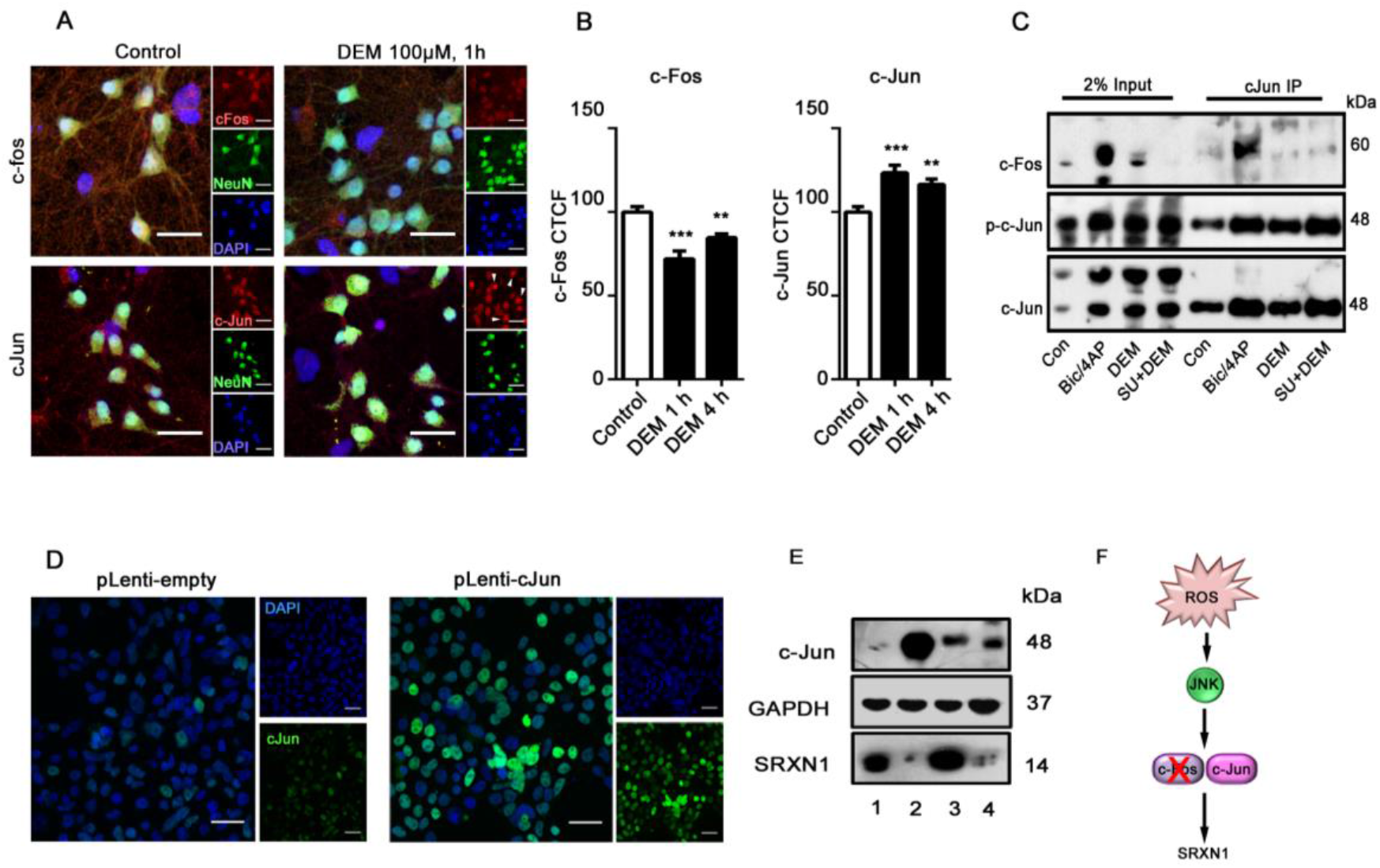
JNK Recruits c-Jun for SRXN-1 expression. ***A+B*** Oxidative stress reduces c-Fos and increases c-Jun. Representative images showing c-Fos (top panels) and c-Jun (bottom panels) immunoreactivity in neurons left untreated (Control) or treated for 1 h with 100 μM DEM. Neurons were co-stained for the neuronal marker NeuN and nuclei were labelled with DAPI. ***B*** Quantification of immunoreactivity in **A**, of neurons stimulated for 1 h or 4 h with 100 μM DEM. Data are plotted as mean ± SEM. Analysis represents NeuN positive neurons. c-Fos levels were assessed in 118 neurons and c-Jun immunoreactivity was quantified in 81 neurons across 3 biological replicates (** p < 0.01; *** p < 0.001 one-way ANOVA followed by a Dunnett’s post-hoc test). Scale bar = 25μm. ***C*** Representative western blot of neuronal lysates immunoprecipitated with a c-Jun antibody in conditions of synaptic activity (Bic/4AP) or oxidative stress (DEM 100 μM) in the presence or absence the JNK inhibitor SU 3327 (700nM, 1 h). ***D*** Representative images of c-Jun immunofluorescence in HEK293T cells transduced with lentivirus encoding empty vector, or pLenti c-Jun. Scale bar = 25μm. ***E*** Western blot showing c-Jun, GAPDH and SRXN-1 levels in neurons transduced with lentivirus containing empty vector (1) or c-Jun (3) overnight, or treated with the Nrf2 agonist, tert-Butylhydroquinone (TBHQ, 10μM, 4) overnight. HEK293T cells transduced with pLenti c-Jun (2) were run as a positive control. ***F*** A schematic demonstrating JNK and c-Jun dependent regulation of SRXN-1 during oxidative stress.

### SRXN-1 is localized at synaptic terminals and prevents dendrite loss

To understand how SRXN-1 contributes to neuronal antioxidant defenses we investigated SRXN-1 localisation in neurons. Having previously identified that chronic treatment of cortical neurons with DEM causes a loss of dendrites, we transfected neurons with PSD95-GFP alone or in combination with a plasmid encoding N-terminal FLAG-tagged human SRXN-1 and treated cells with DEM (48 hours). We found that overexpression of SRXN-1 prevented dendrite loss caused by long-term treatment of DEM (Fig 4A). Interestingly, we also observed overexpressed SRXN-1 immunoreactivity in distal dendrites and on closer examination found that overexpressed SRXN-1 was localised to dendrites and dendritic spines (Fig 4B). SRXN-1 co-localised with PSD-95-GFP, as indicated by white arrowheads in Figure 4B.

**Figure 4:**
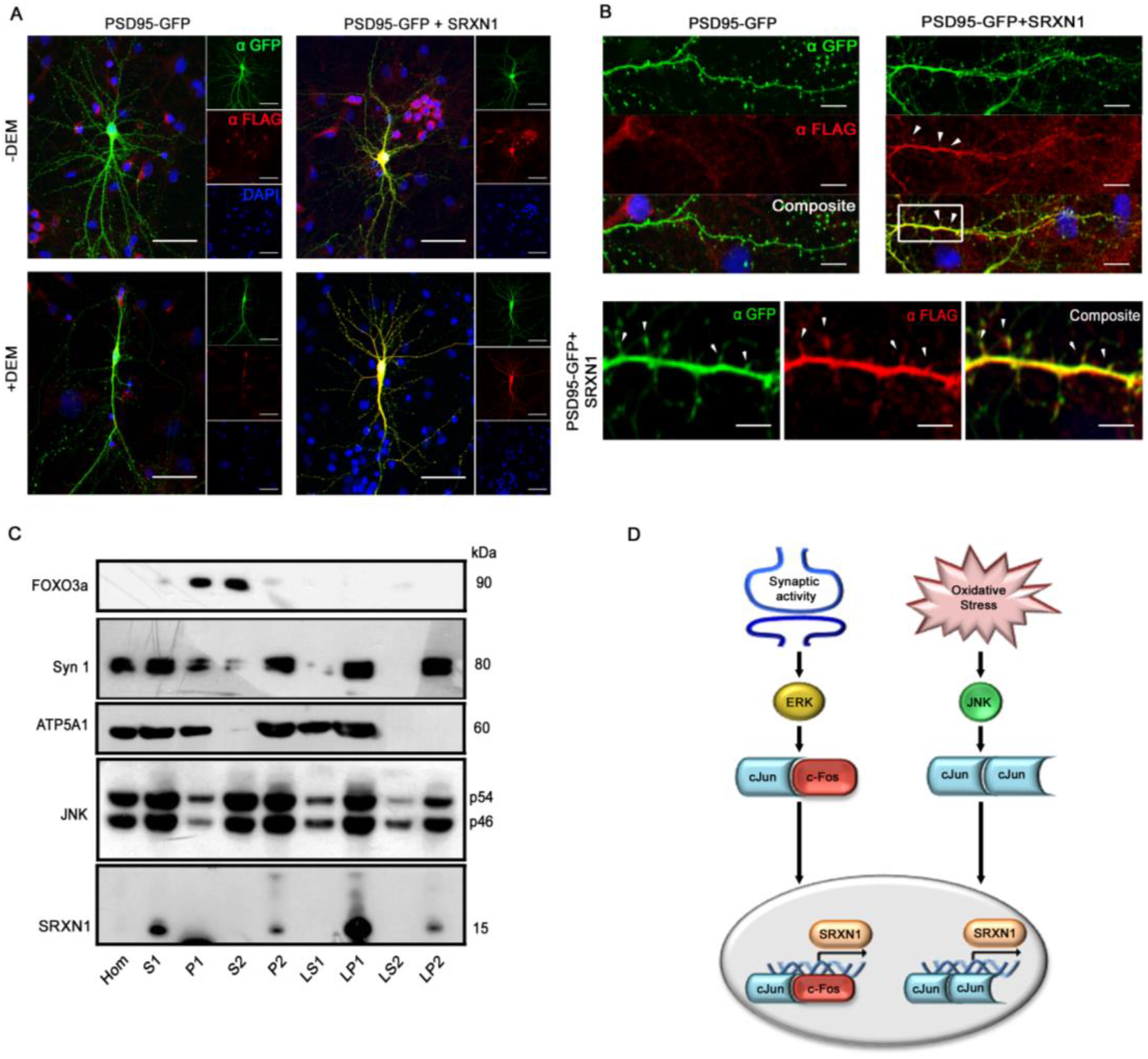
SRXN-1 localises to synaptic terminals in rodent neurons and rescues DEM induced retraction. ***A*** SRXN-1 overexpression rescues DEM-induced dendritic retraction. Representative micrographs of mature neurons transfected with PSD95-GFP constructs alone (left panels) or in combination with Flag-tagged human SRXN-1 (right panels) ± 100μM DEM (48hr). Cells stained with anti-GFP (green), anti SRXN-1, (Flag antibody, red) and nuclear staining with DAPI (blue). Scale bar = 100 μm. ***B*** Overexpressed human SRXN-1 co-localises with PSD95-GFP in dendritic spines. Representative images show an overlay of GFP and Flag immunofluorescence. Magnified images of dendritic spines outlined in white boxes (PSD95-GFP+SRXN-1, scale bar = 5 μm). ***C*** Endogenous SRXN-1 protein is present at synapses. Representative western blot of synaptosome fractions prepared from mouse brain probed with antibodies for the nuclear protein FOXO3a, the pre-synaptic marker synapsin 1 (Syn1), the mitochondrial protein ATP5A1, JNK and SRXN-1 as indicated. ***D*** Schematic representation of likely signaling pathways mediating *Srxn-1* transcriptional induction in response to synaptic activity and oxidative stress.

To further investigate the synaptic localization of SRXN-1, we assessed its distribution in subcellular fractions of mouse brain using the antibody validated in heterologous cells (Fig S4, A). Although characterized as a cytoplasmic enzyme [27], SRXN-1 was not observed in the synaptic supernatant fractions, but in the LP1 (synaptosomal) and LP2 (synaptic vesicle) membrane fractions. The efficacy of the fractionation was confirmed by enrichment of the transcription factor FOXO3a in supernatant fractions over the pellet fractions. The P2 fraction represents a crude synaptosomal fraction comprising microsomes and isolated pre- and post-synaptic nerve terminals. SRXN-1 was most enriched in the LP1 fraction, indicative of synaptic membranes and small organelles, such as mitochondria. Immunoblotting for the alpha subunit of the mitochondrial F1 ATP synthase (ATP5A1) confirmed the presence of mitochondrial proteins in the LP1 fraction. SRXN-1 immunoreactivity was also present in the LP2 fraction, which is enriched in synaptic vesicles (SV), as shown by an enrichment of the SV protein synapsin I (Fig 4C). JNK immunoreactivity was observed in all fractions, but was enriched in P2, LP1 and LP2, in-line with the distribution of SRXN-1. These data suggest that SRXN-1 associates with synaptic membranes and is consistent with the co-localization with PSD95 we observed by fluorescence microscopy. Therefore, in neurons, antioxidant proteins such as SRXN-1 and sensors of oxidative stress such as JNK are located at synapses, placing them at functionally important sites to orchestrate the adaptive antioxidant response (Fig 4D).

## Discussion

Here we have uncovered a role for JNK signalling in the regulation of neuronal antioxidant capacity using *Drosophila* and mammalian neurons in culture. We and others have shown previously that JNK regulates changes in synaptic morphology independently of any stimuli [11] or in response to ROS [5, 12, 14]. We now show that JNK activation drives an antioxidant response, which shapes cellular reductive capacity and synaptic morphology.

We show that *Drosophila* mutants that have enhanced JNK activity have low levels of ROS and furthermore, are resistant to chemically induced oxidative stress. These reduced ROS levels in mutant flies can be reinstated back to levels observed in wild type flies by inhibiting the JNK-AP-1 pathway. In addition to changes in ROS levels, we show that activation of the JNK-AP-1 pathway leads to overgrown synapses at the *Drosophila* NMJ suggesting that the status of JNK signalling has implications for structural synaptic plasticity. In mammalian cortical neurons we identified an ‘adaptive’ JNK-dependent antioxidant response when neurons are challenged with oxidative stress. This adaptive response relies on JNK’s downstream effector c-Jun, and results in increased abundance of the antioxidant protein SRXN-1. In contrast, JNK-independent SRXN-1 expression is triggered with increased synaptic activity of cortical neurons. We therefore implicate JNK signaling in the regulation of neuronal antioxidant defenses and identify distinct differences in neuronal responses to ROS generated in conditions of synaptic activity and oxidative stress.

### Constitutive JNK activation drives an antioxidant response in *Drosophila*

Genetic and pharmacological activation of JNK changes neuronal morphology and this structural change is dependent on AP-1 [15, 28]. JNK-AP-1 signalling has long been known to positively regulate growth and strength at the larval NMJ [15]. JNK has also been shown to mediate the effect of ROS generated during oxidative stress associated with excitotoxicity [14] and lysosomal storage disease [12]. Combined with reports that activation of the JNK pathway after neuronal injury coordinates a regenerative response [16] and that AP-1 signalling regulates dendrite growth during both development and during conditions of synaptic activity; JNK-AP-1 signalling has been demonstrated as a crucial regulator of neuronal homeostasis. Both *puc^E69^*/+ heterozygotes and *hiw* mutants, which have increased JNK activity, [11, 19] showed reduced total ROS levels and rendered flies resistant to chemically induced oxidative stress. The reduced ROS levels in *puc^E69^*/+ heterozygotes and *hiw* mutants flies could be reversed by inhibition of neuronal JNK-AP-1 suggesting that the altered ROS originates from neurons. The reduced total ROS levels, arising from protective JNK signalling, in *hiw* mutants might explain the recently reported resistance of neurons in highwire null flies to the damaging effects of physical blows, which mimic traumatic brain injury [29].

A key question raised by our work is whether the JNK-dependent changes in synaptic morphology are driven by the increased antioxidant defenses. We have previously identified DEM as a regulator of neuronal morphology at the NMJ [5] and here we show that this DEM-induced change in synaptic morphology is regulated by JNK. Furthermore, we show that *puc^E69^*/+ larvae have overgrown NMJ’s which are not influenced by oxidative stress induced by DEM. Given that *puc^E69^*/+ and *hiw* larvae have elevated JNK activity, reduced ROS, elaborate NMJ’s and are resistant to chemically induced oxidative stress, our data suggests that changes in neuronal plasticity in these models is at least, in part, driven by an antioxidant response.

### JNK coordinates antioxidant responses in mammalian neurons

It is known that synaptic activity can regulate the expression of neuronal antioxidant genes [26]. Furthermore, we have recently shown that ROS generated from synaptic activity acts as a signaling molecule to regulate neuronal plasticity, in a PI3K/DJ-1β dependent manner [5] implicating synaptic activity and ROS in the regulation of synaptic plasticity. What is currently unknown is how neurons recognize and respond to differences in physiological and pathological ROS. Our data shows that neurons recruit different signaling pathways to mediate ROS generated by synaptic activity or oxidative stress, but converge to regulate the same antioxidant gene, *Srxn-1*.

Increasing synaptic activity activates ERK1/2 [30, 31] that can in turn phosphorylate AP-1 components [32, 33]. Consistent with this, we found that increasing synaptic activity using Bicuculline and 4-AP regulates *Srxn-1* mRNA and protein expression [26]. Conversely, inducing oxidative stress in neurons with DEM did not affect ERK phosphorylation of AP-1 but did activate JNK and SRXN-1 expression. JNK inhibition abolished increased SRXN-1 in conditions of oxidative stress but not during synaptic activity. This suggests that neurons use distinct kinases to upregulate SRXN-1 expression in response to synaptic activity or oxidative stress. JNK activity can be regulated by cellular glutathione levels. In non-stressed cells, JNK is tethered to monomeric glutathione S-transferase Pi (GST-Pi) and increased H_2_O_2_ can trigger the detachment and oligomerization of GST-Pi, conjugating ROS to glutathione [34]. This suggests that JNK activity is tuned to the redox thresholds of the cell [35]. Both ERK1/2 and JNK kinases regulate AP-1 [21, 32], raising the possibility that the mechanisms converge at the level of this dimeric transcription factor. The AP-1 heterodimer of c-Fos and c-Jun [36] is well characterized but the conditions driving differential heterodimer composition are not well understood. In our work, a notable difference between synaptic activity and oxidative stress conditions was the lack of c-Fos transcriptional/translational induction by DEM and reduced dimerization with c-Jun. This indicates that DEM-induced AP-1 activity and *Srxn1* transcription is c-Fos independent (Fig 4 C+D). The mammalian AP-1 complex can contain Fos, Jun, Fra, and ATF components and multiple hetero- and homo-dimeric partners are capable of binding to AP-1 DNA binding sites [37, 38]. Our immunoprecipitation of c-Jun showed a clear association with c-Fos during conditions of increased synaptic activity, but this association was absent during conditions of oxidative stress. Moreover, total c-Jun increased during oxidative stress and we did not detect JunB or JunD in immunoprecipitates suggesting that c-Jun may homodimerise to generate an antioxidant response in neurons. Our data identify a clear divergence between physiological and stress-dependent signaling in neurons, where activation of JNK and lack of c-Fos defines the stress response.

### SRXN-1 is present at the synapse

SRXN-1 is an oxidoreductase that reduces sulfinylated proteins such as 2-cys peroxiredoxins (PRDX), small redox sensitive proteins widely expressed in mammalian neurons [39]. Both cysteine residues in PRDX can undergo reversible hyper-oxidation by peroxides to form sulfinic acid (Cys-SO_2_H) [40]. Hyper-oxidation of PRDX proteins inactivates their peroxidase activity. This inactivation is known to occur in neurons where it can be seen to follow a circadian time course [41], and occur during glutamate induced excitotoxicity [42] in addition to oxidative stress conditions [43]. SRXN-1 catalyses the reduction of Cys-SO_2_H in an ATP dependent manner, restoring PRDX activity [44]. Both PRDX and SRXN-1 are regulated by synaptic activity [26]. Our data indicate that the cellular levels of SRXN-1 can also be tuned by the intracellular concentration of GSH and H_2_O_2_ levels.

We additionally show that SRXN-1 can be found at cellular locations that are particularly vulnerable to ROS - synaptic terminals and dendrites [45]. Overexpressed human SRXN-1 localised to dendrites and dendritic spines and endogenous SRXN-1 protein was particularly enriched in synaptosomal membrane fractions. We also found that JNK kinases were present in synaptosomes. The localization of SRXN-1, a cytosolic protein, in a synaptic membrane fraction is intriguing. However, in non-neuronal cells, SRXN-1 has been reported to translocate from the cytosol to a ‘heavy membrane’ fraction enriched in mitochondria in response to oxidative stress [46]. Mitochondrial SRXN-1 has a role in re-activation of oxidized peroxiredoxins [40, 47] and our data indicate that SRXN-1 might reside constitutively with neuronal mitochondria, consistent with a greater basal ROS burden in neurons.

Taken together, our data from both fly and mammalian models identify a fundamental conserved role of JNK signaling in regulating neuronal redox homeostasis through an adaptive antioxidant response. JNK signaling is activated specifically by oxidative stress and activates c-Jun containing AP-1 transcription factors independently of c-Fos. In this manner, JNK imparts mechanistic specificity to activate AP-1 under conditions that have relevance to a number of neurodegenerative diseases.

## Acknowledgments

This work was supported by BBSRC Project Grants (BB/I012273/1) awarded to STS and (BB/M002322/1) awarded to STS and SC, and was part-funded by The Wellcome Trust [204829] through the Centre for Future Health (CFH) at the University of York. We thank Addgene (Cambridge, MA. USA.) for providing plasmid stocks. We thank Lucy Rudd for technical assistance and the Bioscience Technology Facility at the University of York for providing access to confocal microscopes.

## Author contributions

C.U., S.C., and S.T.S. designed research; C.U., N.G., S.C., and L.F. performed research; G.E.J. contributed reagents/analytic tools; C.U., S.C., and L.F. analyzed data; and C.U., S.C., and S.T.S. wrote the paper.

## Materials and Methods

### Culture of neurons and cell lines

Timed-mated female Wistar rats (Charles River UK) (RRID:RGD_737929) were maintained in accordance with the UK Animals (Scientific Procedures) Act (1986). Cortices were dissected from postnatal day 1 (P1) mixed sex rat pups. Animals were euthanised using pentobarbital injection followed by cervical dislocation, according to Home Office guidelines. Cortical cell suspensions were obtained as previously described and cytosine arabinoside (AraC, 2.4μM final concentration) was added to the growth medium at 1 DIV [48].

Neurons were transfected at 12 days *in vitro* (DIV) with PSD95-GFP (a kind gift from David Bredt [49] and FLAG-tagged Human SRXN-1 (purchased from Origene: RC207654) for 5 h using Lipofectamine 2000 (11668019, Thermo Scientific). Experiments were performed in a defined culture medium (Transfection medium (TM) contains: 90 ml SGG; 114 mM NaCl, 26.1 mM NaHCO_3_, 5.3 mM KCl, 1 mM MgCl_2_, 2 mM CaCl_2_, 10 mM HEPES, 1 mM glycine, 30mM glucose, 0.5 mM C_3_H_3_NaO_3_ and 10 ml MEM (51200046, Thermo Scientific) supplemented with 100x ITS (1x final concentration; insulin/transferrin/selenium, 41400045, Thermo Scientific), penicillin (50 U/ml) and streptomycin (50 μg/ml)). After 24 h incubation in TM, cells were treated with DEM, bicuculline, catalase, paraquat, rotenone, 4-Aminopyridine or SU3327 at concentrations and durations stated in the figure legends.

HEK293-FT (RRID:CVCL_6911, Thermo Scientific) cells were grown in DMEM containing 10% FBS. Cells were passaged and at 60% confluency, transfected with 10 μg of either pCMV entry vector (empty) or pCMV Human SRXN-1. After 3 days, cells were lysed as described below.

### Lentiviral preparation

Lentiviruses were prepared using the 2nd generation system. Packaging plasmids were produced by the Trono lab (psPAX2 and pMD2.G were gifts from Didier Trono (Addgene plasmid #12260; http://n2t.net/addgene:12260; RRID:Addgene_12260 *and* Addgene plasmid #12259; http://n2t.net/addgene:12259; RRID:Addgene_12259). pLenti c-Jun was created by cloning human c-Jun from PMIEG3-c-Jun (a gift from Alexander Dent (Addgene plasmid #40348)[50] using BAMHI and XHOI into pLenti-puro (a gift from Ie-Ming Shih (Addgene plasmid #39481; http://n2t.net/addgene:39481; RRID:Addgene_39481))[51]. Viral particles were generated and transduced as described previously [52].

### Hydrogen Peroxide Assay

Experiments were performed in phenol red free TM. TM containing 50 μM Amplex Red reagent and 0.1 U/ml HRP was added to neurons in 12 well dishes (500μl per well) for 15 min after which cells were treated with different concentrations of DEM, or catalase. Fluorescence at 590 nm was measured periodically and compared against a H_2_O_2_ standard curve for quantification. For fly experiments, Amplex red reagent was made up in Hemolymph solution (HL3) containing NaCl (128mM), KCl (2mM), CaCl_2_.2H_2_O (1.8mM) MgCl_2_.6H_2_O (4mM), HEPES (5mM) and sucrose (35.5mM) at pH 7.2. Each fly was crushed in 250μl of Amplex red reagent using a pestle, vortexed and left at 25°C, in the dark for 90 minutes. After 90 minutes, all samples were spun down for one minute and each biological replicate was read in duplicate. Only males were used for amplex red assays to avoid issues with dosage compensation.

### WST-1 Assay

After treatment, primary rat neurons were washed with TM and then incubated with phenol red free TM containing WST-1 reagent (Sigma, Cat# 5015944001, 20 μl/ml) for 4 h at 37°C. 200 μl was transferred to a 96 well plate and absorbance measured using a plate reader (BMG Fluostar λ = 440nm).

### Immunocytochemistry

Cells were washed with phosphate buffered saline (PBS) and fixed for 30 min at room temperature with 4% paraformaldehyde (containing 4% sucrose; Sigma). Cells were permeabilized in 0.5% NP40 in PBS for 5 min at room temperature. All primary and secondary antibodies, dilutions and suppliers can be found in Table 1. Primary antibodies were incubated overnight at 4°C. Corresponding Alexafluor secondary antibodies (1:500, Thermo Scientific) were incubated for 1 h at room temperature before mounting with Fluoromount (Sigma).

### Microscopy and Image Analysis

Images were collected on an inverted Zeiss microscope (880) with 20x or 63x Plan Neofluar objectives using Zeiss filter sets for DAPI and Alexa 488, 546 or 633. Images were taken at a resolution of 2048×2048 pixels.

### Synaptosome Preparation

The subcellular fractionation of mouse brain was performed as previously described [53]. Briefly, seven mixed sex mouse forebrains were dissected and transferred into cold homogenization buffer (320 mM sucrose, 1 mM EDTA, 5 mM Tris, pH 7.4, 4°C), homogenized in a glass Teflon homogenizer, and centrifuged in an SS-34 rotor (1000*g*, 10 min, 4°C). The supernatant (S1) was collected and the pellet re-suspended in 15 ml of homogenization buffer and centrifuged again (1000*g* for 10 min). S1 supernatants were combined, and pellets (P1) collected. S1 lysates were further centrifuged at 13000*g* for 20 min at 4°C. Supernatant (S2) was collected and the pellet (P2) re-suspended in 21 ml homogenization buffer. The P2 pellet was further centrifuged at 13000*g* for 20 min at 4°C. P2 was re-suspended in 320 mM sucrose (300 μl/forebrain), transferred into a glass-Teflon homogenizer with 9 volumes of ice-cold distilled water and immediately homogenized for 3 up/down strokes at 2000 rpm. Homogenates were incubated on ice for 30 min with 0.1 volume of 1 M HEPES-NaOH (pH 7.4) and centrifuged at 25000*g* for 20 min at 4°C. Pellets (LP1, synaptosomal membranes) were collected and the supernatant (LS1) further centrifuged in a 70 Ti rotor at 165,000*g* for 2 h at 4°C. The resulting supernatant (LS2) was collected and the pellet (LP2, synaptic vesicles) re-suspended in 40 mM sucrose (2 ml). Protein concentration was assayed using Bradford reagent and 20 μg of each fraction was prepared in 1x Laemmli buffer for western blotting.

### Western Blotting

Cells were lysed in RIPA containing phosSTOP phosphatase inhibitors (4906845001, Roche) and complete EDTA-free protease inhibitors (04693132001, Roche) as previously described [54]. Lysates were run on Novex pre-cast mini gels (NuPAGE 4-12% Bis-Tris Gels, NP0322BOX, Thermo Scientific) in either 1 x MES or 1 x MOPS buffer. Antibodies used for immunoblotting are detailed in Table 1.

### Co-immunoprecipitation

Neurons in 10cm dishes were lysed as described. Lysates were incubated with primary antibodies for c-Jun at a 1:50 dilution and rotated overnight at 4°C. Lysates were incubated with 15μl (30μl bead slurry) Protein G, Sepharose (GE Healthcare, 17-0618-01) for 2 h at 4°C rotating. Samples were spun down, transferred to spin-x columns (Sigma, CLS8163-100EA), washed 3 times with lysis buffer and the beads finally incubated with 4 x Laemmli (containing mercaptoethanol) for 10 minutes, then eluted through the spin columns. Samples were further heated at 95°C for 5 minutes before western blotting. Proteins were detected using conformation specific antibodies (Cell Signaling, L27A9, mAb #3678) which prevent detection of heavy and light antibody chains.

### Glutathione Assay

Total levels of glutathione were assayed using a colorimetric Glutathione Assay Kit (CS0260, Sigma). Briefly, at 14 DIV, primary neurons in 35 mm dishes were treated with DEM in transfection medium for 1, 4 and 24 h. Cells were then washed with PBS (4°C), lysed with 5% sulfosalicylic acid and snap frozen in liquid nitrogen. Lysates were then defrosted at 37°C, centrifuged at 10,000*g* for 10 min and 10 μl samples were used for the glutathione assay in accordance with the manufacturer’s instructions.

### Quantitative PCR

Real-time quantitative PCR on the resulting cDNA was performed using a Fast SYBR Green Master mix (Applied Biosystems, 4385612) and gene specific primers. Relative expression of genes was determined by the 2-ΔΔCT method and Gapdh for normalization. Primers used were: Gapdh forward: 5’-AAACCCATCACCATCTTCCA-3’ and Gapdh reverse: 5’-GTGGTTCACACCCATCACAA-3’; c-Fos forward: 5’-AGAATCCGAAGGGAAAGGAA-3’ and c-Fos reverse: 5’-ATTGAGAAGAGGCAGGGTGA-3’; Srxn1 forward: 5’-GACGTCCTCTGGATCAAAG-3’ and Srxn1 reverse: 5’-GCAGGAATGGTCTCTCTCTG-3’

### *Drosophila* stocks and husbandry

*Drosophila* were raised on 4-24® instant *Drosophila* medium (Carolina Biological Supply Company, USA) supplemented with a yeast sucrose solution (5% w/v inactivated yeast, 10% w/v sucrose in ddH_2_O, 100g/500ml) and maintained at 25°C on a 12 h light:dark cycle. Prior to mixing with 4-24® instant media, vehicle (Ethanol) and DEM were added at the desired concentration (Ethanol 0.16%, DEM 0-10 mM). The following stocks were obtained from Bloomington *Drosophila* stock center: Canton-S (CS), *w^1118^*, UAS-*bsk^K53R^* (#9311), UAS-*fos^DN^* (#7214), UAS-*jun^DN^* (#7217), Act5C-Gal4 (Actin-Gal4), nsyb-Gal4. UAS-*ask1^K618M^* (*ask1^DN^*) was a kind gift from Masayuki Miura [55]. UAS-*wnd^DN^* (*wnd^KD^*) and *hiw^ND9^* were kind gifts from Aaron DiAntonio [11, 56]. *puc^E69^/TM6b* flies were obtained from Alfonso Martinez-Arias [57]. *Spin*Gal4/TM6b was obtained from Daisuke Yamamoto [58]. The JNK dominant negative (*bsk^K53R^*)[59] and the *highwire* mutant (*hiw^ND9^*) have been described previously [18]. All analyses were performed in males. *hiw^ND9^* females were crosses to Canton-S males to produce male *hiw*ND9 animals in an outcrossed background. For *hiw^ND9^* crosses with dominant negative transgenes, *hiw^ND9^*; nsyb-Gal4 females were crossed with males carrying the UAS transgenes. All wild types were an outcross of Canton S to w^1118^.

### Survival Analysis

Hatched 1^st^ instar larvae were collected and raised in standard food with different DEM concentrations (0 mM, 1mM, 5 mM and 10 mM). For each survival experiment, at least 2 vials, each containing 50 larvae, were maintained until fly eclosion. The number of eclosed flies was recorded.

### Immunohistochemistry and NMJ Analysis

Third instar wandering larvae were dissected, fixed, antibody stained, imaged and analysed as described previously [13]. All NMJ analysis was performed double-blind. Primary antibodies detailed in Table 1. Confocal microscopy was performed using a Zeiss LSM 880 on an Axio Observer.Z1 invert confocal microscope (Zeiss). Z-stacked projections of NMJ’s and VNCs were obtained using a Plan Neofluar 40x/0.75 NA oil objective. NMJ lengths were measured from stacked NMJ images using the NeuronJ plugin for ImageJ (National Institutes of Health) as described previously [13, 54].

### Antibodies

| Name | Supplier | Dilution | Species | RRID |
| --- | --- | --- | --- | --- |
| Anti- GFP | Synaptic Systems | 1:1000 | Guinea Pig | RRID:AB_11042617 |
| Anti- Synapsin 1 | MERCK | 1:1000 | Rabbit | RRID:AB_2200400 |
| Anti- GCLC | SCBT | 1:500 | Mouse | RRID:AB_2736837 |
| Anti- GCLM | SCBT | 1:500 | Mouse | RRID:AB_831789 |
| Anti- Phospho JNK | Cell Signalling Technology | 1:1000 (WB) | Mouse | RRID:AB_2307321 |
| Anti SAPK/JNK Antibody | Cell Signalling Technology | 1:1000 (WB) | Rabbit | RRID:AB_2250373 |
| Anti FOXO3a | Cell Signalling Technology | 1:1000 (WB) | Rabbit | RRID:AB_2636990 |
| Anti- FLAG | Sigma | 1:1000 | Mouse | RRID:AB_259529 |
| Anti- FLAG | Cell Signalling Technology | 1:1000 | Rabbit | RRID:AB_2217020 |
| Anti- c-Jun | Cell Signalling Technology | 1:1000 (WB) | Rabbit | RRID:AB_2130165 |
| Anti- Phospho c-Jun | Cell Signalling Technology | 1:1000 (WB) | Rabbit | RRID:AB_2130162 |
| Anti- NeuN | Millipore | 1:2000 | Guinea Pig | RRID:AB_11205592 |
| Anti- c-Fos | Cell Signalling Technology | 1:2000 (WB) | Rabbit | RRID:AB_2106617 |
| Anti- GAPDH | Millipore | 1:10,000 (WB) | Mouse | RRID:AB_2107445 |
| Anti- JunB | SCBT | 1:1000 (WB) | Mouse | RRID:AB_2130023 |
| Anti- JunD | SCBT | 1:1000 (WB) | Mouse | RRID:AB_10650101 |
| Anti- SRXN-1 | SCBT | 1:250 (WB) | Mouse | RRID:AB_2286615 |
| Anti- ATP5A1 | Proteintech | 1:1000 (WB) | Rabbit | RRID:AB_2061761 |
| Anti- p44/42 MAPK (Erk1/2) | Cell Signalling Technology | 1:1000 (WB) | Rabbit | RRID:AB_330744 |
| Anti- Phospho-p44/42 MAPK (Erk1/2) | Cell Signalling Technology | 1:1000 (WB) | Mouse | RRID:AB_331768 |
| **Drosophila NMJ Antibodies** |  |  |  |  |
| Anti- Horseradish peroxidase- Cy3 (HRP-Cy3) | Jackson ImmunoResearch Labs | 1:200 | Goat | RRID:AB_2307391 |
| Anti-synaptotagmin (Anti-SYT91) | Sweeney Lab | 1:2000 | Rabbit | RRID:AB_2713991 |
| **Secondary Ab's** |  |  |  |  |
| Gt anti Ms Alexa 488 | Life Technologies | 1:500 | Goat | RRID:AB_2576217 |
| Gt anti Rb Alexa 546 | Life Technologies | 1:500 | Goat | RRID:AB_10563566 |
| Gt anti Ms Alexa 546 | Life Technologies | 1:500 | Goat | RRID:AB_2534071 |
| Gt anti Rb Alexa 488 | Life Technologies | 1:500 | Goat | RRID:AB_2576217 |
| Peroxidase-AffiniPure Goat Anti-Guinea Pig IgG (H+L) antibody | Jackson ImmunoResearch Labs | 1:10,000 (WB) | Goat | RRID:AB_2337402 |
| Peroxidase-AffiniPure Goat Anti-Rabbit IgG | Jackson ImmunoResearch Labs | 1:10,000 (WB) | Goat | RRID:AB_2307391 |
| Peroxidase-AffiniPure Goat Anti-Mouse IgG (H + L) antibody | Jackson ImmunoResearch Labs | 1:10,000 (WB) | Goat | RRID:AB_10015289 |
| Fluorescein (FITC) AffiniPure Goat Anti-Rabbit IgG (H+L) | Jackson ImmunoResearch Labs | 1:200 | Goat | RRID:AB_2337972 |
| **IP Ab's** |  |  |  |  |
| Mouse Anti-rabbit IgG (Conformation Specific) (L27A9) mAb #3678 | Cell Signalling Technology | 1:5000 (WB) | Mouse | RRID:AB_1549606 |
| Anti-mouse IgG, HRP-linked Antibody #7076 | Cell Signalling Technology | 1:10,000 (WB) | Horse | RRID:AB_330924 |

## Supplementary Figure Legends

**Figure S1.**
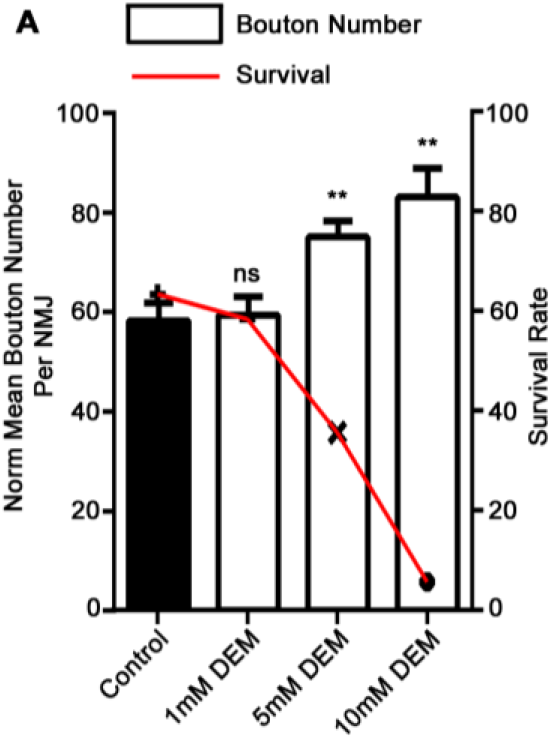
Effects of DEM on NMJ bouton number and survival in *Drosophila*. DEM (0 mM, 1 mM, 5 mM and 10 mM) was added to standard food and NMJ bouton number quantified. NMJ bouton number increases as DEM concentrations increase. (left y-axis, bars). Survival significantly decreases as DEM concentrations increase (right y-axis, red line). Data plotted as means ± SEM analysed using one-way ANOVA with a Dunnett’s post hoc (** p < 0.01).

**Fig S2:**
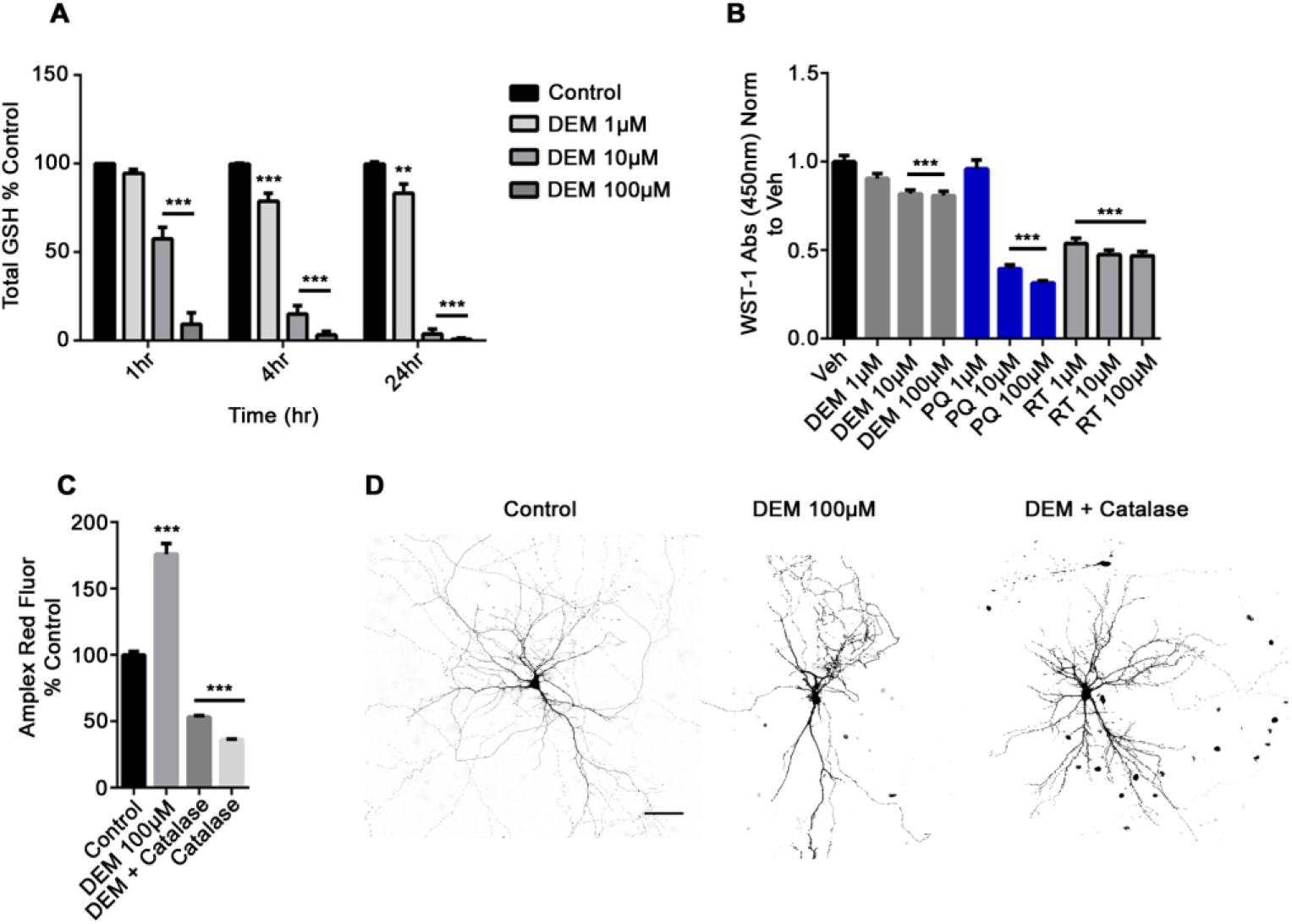
DEM depletes glutathione and induces oxidative stress. ***A***. Total glutathione content in mature neurons treated with DEM at the indicated time and concentration. Total glutathione content is relative to ethanol treated controls and data are shown as means ± SEM (** p < 0.01; *** p < 0.001, using two-way ANOVA followed by a Dunnett’s *post-hoc* test; *n* = 3 biological replicates). ***B*** Mitochondrial toxicity was assessed using WST-1 assays in neurons treated with DEM, paraquat (PQ) and rotenone (RT) at the indicated concentrations, for 24 hours. WST-1 turnover is normalized to ethanol treated controls (Veh) and shown as means ± SEM (***p < 0.001; one-way ANOVA; n = 3 biological replicates). ***C*** H_2_O_2_ production was monitored using Amplex red reagent. Neurons were treated with DEM and fluorescence measured after 24 hours. Catalase (100 nM) was added to neuronal cultures 15 min before DEM treatment. Data represents means ± SEM analysed using one-way ANOVA (*** p < 0.001, n = 3 biological replicates). Amplex red fluorescence is shown as a percentage of vehicle (0.1% ethanol) treated controls. ***D*** Prolonged oxidative stress causes dendritic loss. Representative micrographs of neurons transfected with PSD95-GFP and treated, with either 0.1 % ethanol (control) or the indicated concentrations of DEM (100μM) +/− Catalase (100nM) for 48 h. Scale bar = 50 μm.

**Fig S3.**
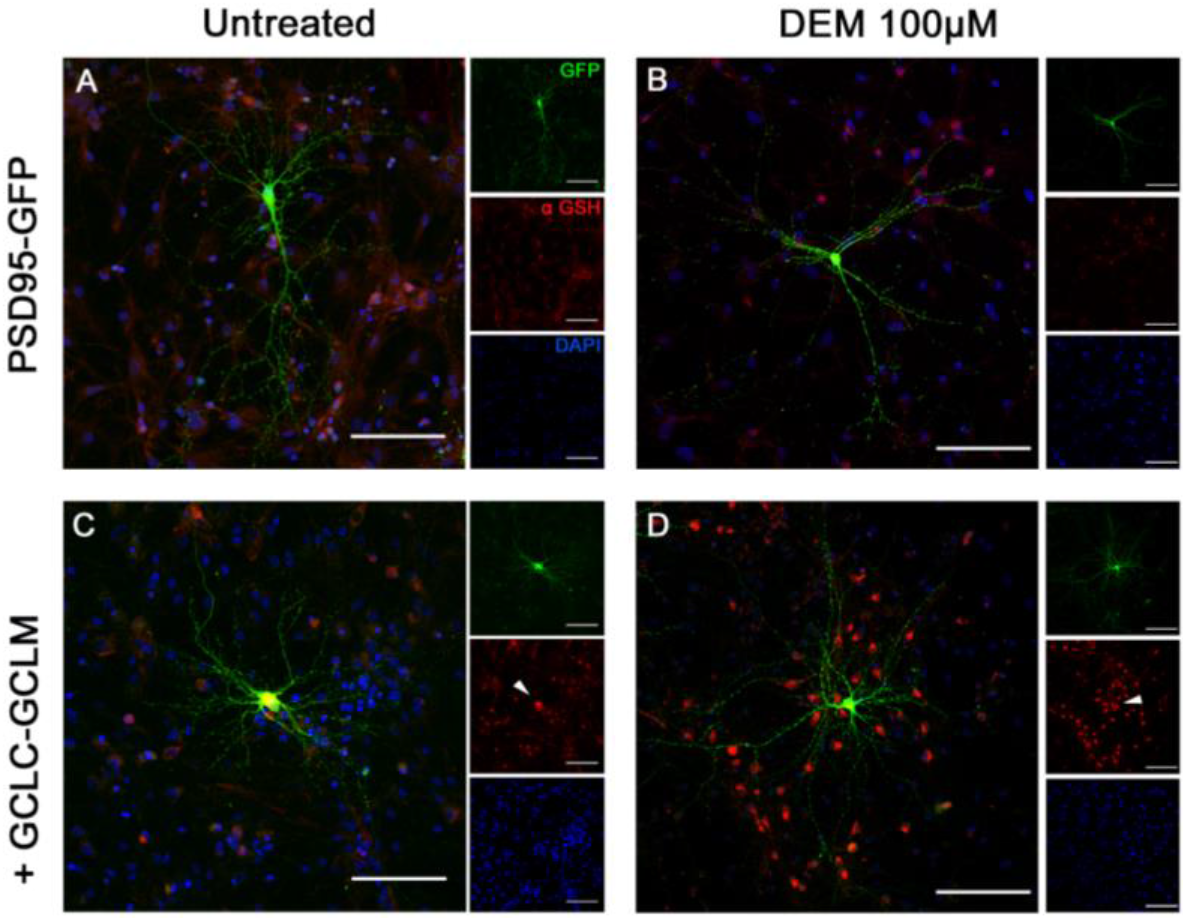
DEM induced dendritic loss is rescued by GSH overexpression. **A-D** Mature neurons transfected either with a PSD95-GFP expression plasmid alone (top panels) or PSD-95GFP along with plasmids expressing the catalytic (GCLC) and modifying (GCLM) subunits of Glutamate Cysteine Ligase (bottom panels). Cells were incubated with either vehicle (0.1% ethanol) or DEM (100 μM) for 48 h (scale bar = 100 μm). Insets show anti-GFP (green), anti-GCLC (red) and nuclei (blue).

**Fig S4.**
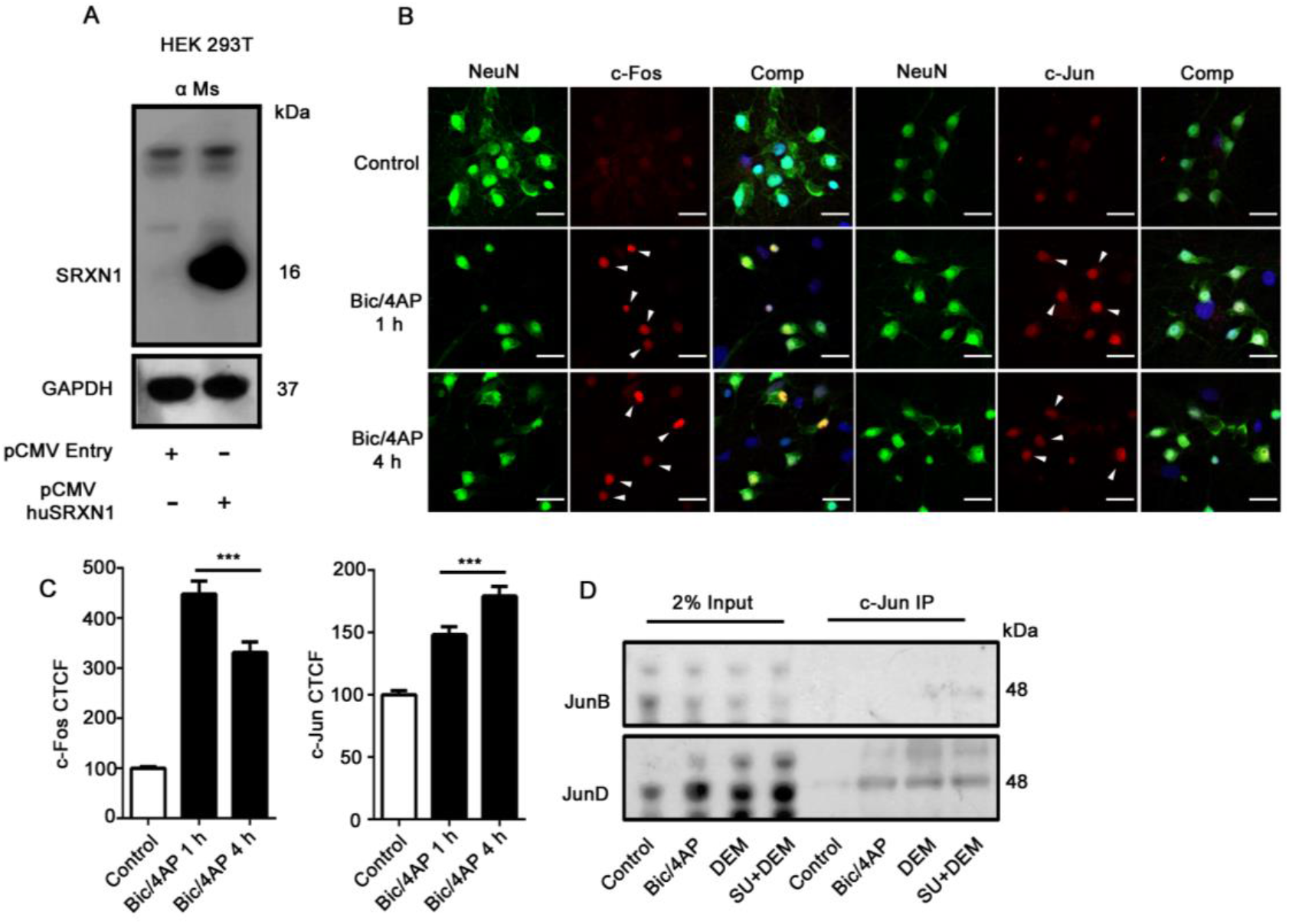
Synaptic activity induced c-Fos and c-Jun. **A** Representative western blot of protein samples from HEK293T cells transfected with a FLAG-tagged Human SRXN-1 expression vector and probed with anti-SRXN-1 mouse monoclonal antibody (top panel, SI Appendix, Table 1) or GAPDH antibody (bottom panel). **B** Synaptic activity induces c-Fos and c-Jun. Representative images showing c-Fos and c-Jun immunoreactivity in neurons left untreated (Control) or treated for 1 h or 4 h with Bicuculine (50 μM, Bic) and 4-Aminopyridine (500 μM, 4AP). Neurons were co-stained for the neuronal marker NeuN and nuclei were labelled with DAPI. Scale bar = 20μm. **C** Graphs showing c-Fos (left) and c-Jun (right) levels in neurons treated for 1 h or 4 h with Bic/4AP. Data are plotted as mean ± SEM. Analysis represents NeuN positive neurons and c-Fos levels were assessed in 135 neurons across 3 biological replicates (*** p < 0.001 one-way ANOVA followed by a Dunnett’s post-hoc test). c-Jun immunofluorescence was assessed in 78 neurons across 3 biological replicates (*** p < 0.001 one-way ANOVA followed by a Dunnett’s post-hoc test). CTCF = corrected total cell fluorescence analysis. **D** Representative immunoprecipitation of c-Jun in conditions of increased synaptic activity (Bic/4AP, 1 h) and oxidative stress (DEM, 100 μM, 1 h) and western blot showing levels of JunB (upper panel) and JunD (lower panel).

